# Species-level variation in primate social behavior is correlated with climate extremes and variability

**DOI:** 10.64898/2026.06.15.732406

**Authors:** Maria J.A. Creighton

**Affiliations:** Department of Biology, Duke University, Durham, NC, USA

**Keywords:** climate, group size, primates, social grooming, social behavior

## Abstract

Cooperatively breeding species are disproportionately found in extreme and unpredictable climates globally, suggesting that cooperation is beneficial to persistence in climatically challenging conditions. Notably, other dimensions of sociality, like group living and tendency to engage in affiliative social behaviors, offer fitness-related benefits that could make them similarly advantageous in such climates. Here, I present a phylogenetic analysis of Primates aimed at testing whether these two dimensions of sociality—average group size and average percent time spent social grooming—are predicted by climatic challenges in species’ environments. Results show that time spent grooming is highest in extreme and unpredictable climates, with how dry conditions are explaining the greatest amount of variation. Thus, climate may influence the evolution and/or persistence of social grooming. While multiple mechanisms could mediate this association, subsequent analyses point to the benefits of social affiliation in environments where groupmates have highly competitive dynamics as one potential explanation.

## INTRODUCTION

The tendency to be social is thought to be favored in challenging environments. For instance, theoretical models predict that cooperation should be selected for when external conditions are harsh or uncertain (Andras et al. 2007; Smaldino et al. 2013; Komdeur & Ma 2021). Many potential sources of environmental harshness and uncertainty for wild animals are a product of climatic conditions. Variation in key climatic variables like rainfall and temperature determine resource availability (reviewed in White 2008) and the physiological demands of thermoregulation (reviewed in Tattersall et al. 2012). Thus, the tendency for animals to be social may be selected for in climates where harshness and uncertainty in rainfall and temperature create adverse conditions.

In line with this prediction, analyses show that one species-level social trait, the tendency to cooperatively breed, is correlated with climatic variables: cooperatively breeding birds and mammals are more likely to be geographically located in climates characterized by extremes or uncertainty in rainfall and temperature, than non-cooperative species (Rubenstein & Lovette 2007; Jetz & Rubenstein 2011; Cornwallis et al. 2017; Lukas & Clutton-Brock 2017). Evidence from ancestral state reconstructions in birds suggests that this pattern is a result of cooperation promoting the invasion of challenging environments, as opposed to cooperative breeding evolving in challenging conditions (Cornwallis et al. 2017). Regardless of whether cooperative breeding evolves in challenging environments or improves invasion propensity into challenging environments, these cross-species analyses reveal a potential advantage of cooperative breeding for survival and persistence challenging climates.

Importantly, the tendency to breed cooperatively represents only one dimension of animal sociality. Animal species vary considerably in other dimensions of sociality that are also associated with fitness and survival outcomes (Silk 2007) and may thus also offer advantages in challenging climates. For instance, living in large groups can improve competitive ability and ultimately foraging success (Hamilton et al. 1975; Packer et al. 1990; Creel & Creel 1995), which may be particularly important when resources are limited or unpredictable. Having a larger number of social partners can also offer thermoregulatory benefits, which may be most beneficial in extreme temperatures (Majolo et al. 2013; McFarland & Majolo 2013; Russo et al. 2017; Campbell et al. 2018). Meanwhile, affiliative social behaviors like allogrooming (hereafter referred to as social grooming) can offer benefits that include reducing stress during times of adversity (Taylor et al. 2003; Wittig et al. 2008), reducing external parasites (Brooke 1985; Akinyi et al. 2013; Villa et al. 2016; Russo et al. 2017), and helping with thermoregulation (McFarland et al. 2016), all of which may be particularly important for survival in challenging climates. Affiliative behaviors can also act as currency in competitive landscapes (e.g., in exchange for social support when there is high within-group competition). A greater expression of these behaviors may therefore be favored in extreme or uncertain climates if these conditions create a patchier distribution of resources and more within-group competition (Henzi & Barrett 2003). Despite their possible role in helping animals to navigate adverse conditions, it is largely unknown whether either of these dimensions of sociality—group size or social grooming—covary with climate.

Primates are one of the most species-rich mammal orders (>500 species; IUCN 2025) with an abundance of data available on their social organization and behavior. Species of non-human primates also engage in the same affiliative behaviors as one another (e.g., social grooming; Seyfarth 1977; Dunbar 1991) allowing for direct cross-species comparisons in the tendency of individuals to socially affiliate. Moreover, non-human primates are widespread, with a global range encompassing ∼46 million km^2^ spanning Africa, Asia, Europe, and the Americas (Estrada et al. 2022), and individual species that tend to be geographically clustered. Thus, unlike globally widespread lineages where species are not geographically clustered, there is distinct species-level variation in the average climatic conditions experienced by different primate species. These facts combined with emerging evidence that climate has played an important role in shaping other aspects of primate social evolution (Kamilar & Baden, 2014; Qi et al. 2023), makes primates an ideal taxonomic group for exploring the relationship between climate and species-level variation in sociality.

Here, I investigate the relationship between the climate conditions in primate species’ environments and two species-level social measures: average group size and average percent time spent social grooming. These two social measures capture species-level variation in primate sociality. I test how each of these measures correlates with an Index of Climatic Challenges (ICC), with the prediction that larger groups and engagement in affiliative social behaviors are more common in challenging climates.

I find that while climatic extremes and uncertainty do not predict group size, the ICC is a strong predictor of the average percentage of time individuals spend grooming across species in the predicted direction: species in more climatically challenging environments spend more time grooming, with the degree to which an environment is dry seemingly having the largest influence in driving this relationship. Multiple non-mutually exclusive mechanisms could cause social grooming to be favored in challenging environments. Social grooming could function to reduce stress in challenging environments, enhance thermoregulation, reflect increased investment in social partners due to higher-within group competition in these climates, or help to combat increased ectoparasite demands. This pattern could also be explained if challenging climates select for uncaptured aspects of social organizations/systems that intensify selection for affiliative behaviors. These mechanisms are difficult to disentangle using empirical data; however, I set out to explore two of them—differences in social organization/systems and ectoparasite richness— using publicly available data compilations and discuss the implications of these results for our understanding of social evolution.

## METHODS

### Data

#### Sociality measures

The full dataset used for this analysis contained 275 species included in the trait data compilation in Creighton & Nunn (2023) that had available published data on group size and/or percent time spent grooming. Data on average group size for species (range: 1 to 242 average group members) were taken from the compilation in Creighton & Nunn (2023), which created averages from compiled sources in Rowe & Myers (2011). Data on the average percentage of time individuals in different species spent social grooming were taken from the data compilation published in Grueter et al. (2013) (range: zero to 18 percent of the average individual activity budget spent social grooming). This data compilation contains behavioral averages across individuals in established study populations, with each species being represented by between one and three representative study populations (Fig. S1).

#### Index of Climatic Challenges (ICC)

To quantify the abiotic challenges in species’ global ranges, species’ range polygons were sourced from the IUCN (2025). Climatic data for each species’ geographic range were then extracted from WorldClim Version 2 (Fick & Hijmans, 2017) using the R package “geodata” (Hijmans et al. 2024). Each species’ geographic range received a mean value (taken across grid cells and weighted by the percentage of each cell that fell in the species’ range) of mean annual temperature (°C), temperature seasonality (defined by WorldClim as the standard deviation in average monthly temperature across the 12 months of the year x 100; Fick & Hijmans, 2017), mean annual precipitation (mm), and precipitation seasonality (defined by WorldClim as the coefficient of variation in long-term average total monthly precipitation x 100; Fick & Hijmans, 2017). These four metrics were then combined into a single standardized index that captured climate extremes and uncertainty, with higher values representing those conditions thought to be most physiologically demanding for primates. To do this, each climate measure was standardized by centering around the mean and scaling within one standard deviation and modified such that higher values indicated more challenging environmental conditions. Specifically, standardized mean precipitation was reflected about the mean, given the demonstrated influence of drought on physiological stress and health in multiple species (Young et al. 2019; Campos et al. 2020; Carrera et al. 2025). Meanwhile, mean temperature was used to calculate each species’ deviation from the mean temperature experienced across species given that both high and low temperature extremes present physiological challenges for primates (McFarland & Majolo 2013; Lucore et al. 2024). Temperature deviation was then standardized to be on the same scale as the other climate variables. Greater climatic seasonality creates more variability in key resources which primates must cope with through behavioral or physiological adaptations (van Schaik & Pfannes 2005), and as such standardized seasonality variables were not modified.

An Index of Climatic Challenges (ICC) was established by summing the four climate variables described above: i) standardized reflected mean precipitation, ii) standardized temperature deviation, iii) standardized temperature seasonality, and iv) standardized precipitation seasonality (Fig. 1). High values of this index thus represent climates characterized by low and heterogeneous rainfall and by extreme and heterogenous temperature. While this index does not come close to capturing all environmental challenges species may endure, it captures similar environmental adversities to those demonstrated to be correlated with cooperative breeding in published analyses, suggesting social traits respond to these climate variables (Rubenstein & Lovette 2007; Jetz & Rubenstein 2011; Cornwallis et al. 2017; Lukas & Clutton-Brock 2017). Notably, the climate index developed for this analysis was not strongly correlated with species’ global range sizes (Fig. S2), suggesting differences in ICC scores were not driven by differences in how globally widespread species were.

**Fig. 1:**
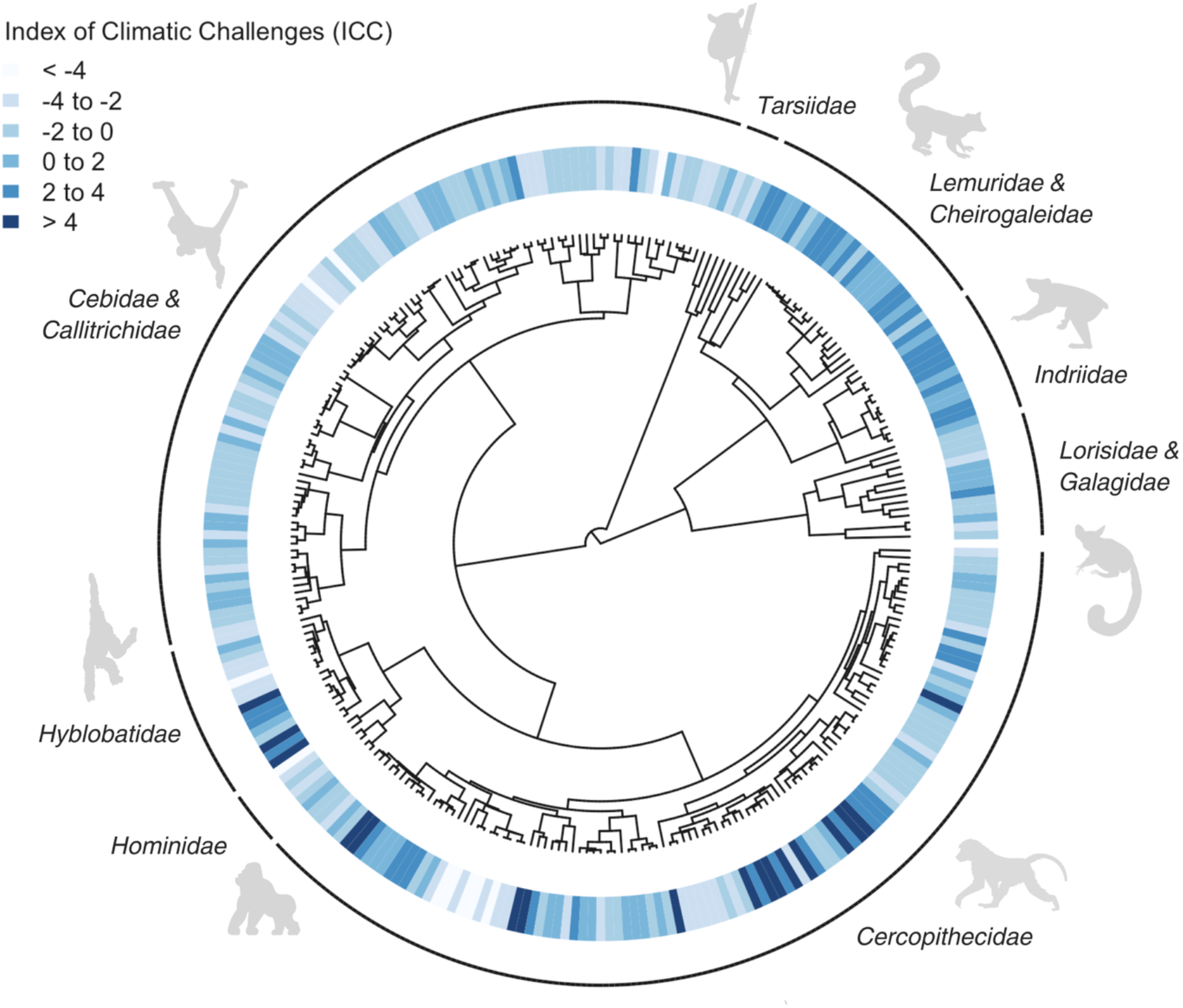
Phylogenetic distribution of how species’ geographic ranges score on an Index of Climatic Challenges (ICC) for 275 primate species plotted on a sampled phylogeny from Upham et al. (2019). Darker colors correspond to more challenging (i.e., more extreme and variable) environments.

#### Other covariates

Differences in life histories or lifestyle that are not social may influence the social tendencies of mammals. To account for this possibility, data on average adult body mass (in grams)—a proxy of many primate life history traits including generation length, development, and lifespan (Peters 1986)—and whether or not species were terrestrial/semi-terrestrial (true=1 and false=0) were taken for primate species from Galán-Acedo et al. (2019) and Rowe & Myers (2011), respectively, and supplemented with data from other sources (see Creighton & Nunn 2023).

#### Phylogeny

A block of 100 maximum likelihood ultrametric mammal phylogenies were selected at random from Upham et al. (2019). These phylogenies were trimmed to only include relevant primate species (i.e., those species with available data) and used to control for phylogenetic relatedness and uncertainty in all analyses.

### Analysis

All data analyses were conducted using R version 4.5.1 (R Core Team (2025).

#### Climate as a predictor of species-level social measures

To test if the ICC predicted species-level average group size or average percent of time spent social grooming, two Bayesian phylogenetic models were built using the “MCMCglmm” package (Hadfield, 2024) in R. Both social measures had relatively high phylogenetic signal (see Supplementary Materials), suggesting a phylogenetic framework was appropriate for these analyses. In each model, one of the two social measures (average group size or average percent time spent grooming) was an outcome, and the ICC was a fixed effect. Average adult body mass (a continuous variable) and whether the species was primarily terrestrial/semi-terrestrial (true=1 and false=0) were included as additional fixed effects to account for life history and lifestyle differences that might account for variation in social measures. Continuous fixed effects were centred with respect to their means and scaled by one standard deviation to improve mixing and in accordance with fixed effects priors. Each model included all species that had data coverage for the response variable and all fixed effects; n=259 species for the average group size model and n=70 species for the average percent time spent grooming model. Limitations on the sample sizes were mostly determined by data coverage of the response variables, although some species with group size data were missing data on body mass, and were thus also excluded.

The model for average group size was assigned a Gaussian distribution and percent time spent grooming was logit-transformed and also assigned a Gaussian distribution. In both models the fixed effects were assigned a weakly informative Gelman prior while the random effect of phylogeny and variance components were assigned weakly informative inverse-Wishart priors. The model with average percent time grooming as an outcome was run a second time using an ICC calculated from study population-specific climate data (as opposed to species’ entire geographic ranges), since few representative populations contributed to species-level grooming values. In cases where species-level grooming data came from averaging data across multiple populations, site-specific ICC scores for these populations were also averaged to get a mean site-specific ICC value for the species.

All models ran a total of 800,001 iterations with a thinning interval of 500 and burn in of 10,000. Models were confirmed to have converged via inspection of trace plots and diagnostics. Each described model was run over all 100 randomly sampled phylogenies from Upham et al. (2019) to account for phylogenetic uncertainty. Results from models run across all 100 phylogenies were averaged to provide the mean effect sizes reported in the text and in the summary tables presented in the Supplementary Materials.

In addition to testing the ICC as a predictor of social measures, temperature deviation, temperature seasonality, reflected mean precipitation, and precipitation seasonality were tested as separate predictors of social measures to assess their relative influence in driving possible correlations. In these analyses, in addition to looking at seasonality within year, temperature and precipitation variability across years were also tested as predictors to ensure the scale of climate variability did not influence results. Data for these analyses came from TerraClimate (Abatzoglou et al. 2018), which were used to calculate inter-annual (1970 to 2025) measures of variability following the same approach used by Fick & Hijmans (2017) to estimate within-year seasonality i.e., the standard deviation x 100 of average annual temperature and the coefficient of variation in total annual precipitation x 100.

#### Social organization and parasite analysis

Several mechanisms that are not mutually exclusive could link challenging climates to the percentage of time individuals from different species spend social grooming, many of which would be difficult to investigate using empirical data. However, existing data compilations allowed for an investigation of two mechanisms—differences in social organization/systems and ectoparasite richness—in mediating a potential relationship.

To explore whether differences in social organization or systems could explain the relationships between social grooming and the ICC, the model for percent time spent grooming, described above, was ran a second time with average group size, social system (solitary, pair living, polygyny, and polygynandry; where the baseline was solitary), and female philopatry (true=1 and false=0) as additional predictors (see Supplementary Methods). This analysis included 62 species that had data available for percent time spent grooming, body mass, terrestriality, and these additional social covariates. I focused on these three aspects of sociality because of their posited relationship to the variables of interest. Specifically, average group size shares a demonstrated positive correlation with social grooming time (Dunbar 1992; Lehmann et al. 2007; Dunbar & Lehmann 2013; Fig. S2; but also see Grueter et al. 2013), social system is thought to influence the amount of time spent with conspecifics and the social functions of grooming (Dunbar 1988), and female philopatry may emerge in challenging climates and lead to more grooming among females (Lehmann et al. 2007; Dunbar & Lehmann 2013).

Differences in dominance systems may also be linked to climate, given that social hierarchies are thought to evolve in response to resource distribution (Boone, 1992) and are known to affect grooming relationships (Seyfarth, 1977; Qi et al. 2023). However, the limited number of species with data in representative cross-species compilations of dominance styles (see Supplementary Methods) meant sample size limitations prevented these metrics from being included as predictors in formal analyses. Instead, two key indicators of dominance style, aggression symmetry (measured using the directional inconsistency index; see Supplementary Methods) and counteraggression (measured as the percent of aggressive bouts followed by retaliation; see Supplementary Methods), were sourced from Kavanagh et al. 2021 for representative species from several families and plotted against both percent time spent grooming and the ICC to provide insights about the potential role of dominance dynamics in mediating a percent time spent grooming-ICC correlation.

To explore whether differences in ectoparasites could explain a link between percent time spent grooming and the ICC, data on ectoparasite-host interactions were sourced from the Global Mammal Parasite Database (GMPD) (Nunn & Altizer, 2005). While these data did not allow for an estimate of the total abundance of ectoparasites in a habitat for which a species is a viable host, as would be ideal for assessing the hygienic demands of grooming, they did allow for a rough estimate of the number of known ectoparasite species that are found on a host taxa. Species richness can reflect the suitability of a given ecosystem in supporting ectoparasites that affect a given host, and species richness is posited to increase with total number of individuals (Srivastava & Lawton, 1998) and may thus reflect a species’ average ectoparasite burden. Nevertheless, considering the limitations of this measure the results from these models should be treated as exploratory. Using counts of the total number of ectoparasite types known to affect each primate species and GMPD sampling effort as additional fixed effects (see Supplementary Methods), models for percent time spent grooming were rerun to see if these data could reveal any effect of ectoparasite differences in explaining the relationship between grooming and the ICC. This analysis included 55 species that had data for both percent time spent grooming and data in the GMPD.

## RESULTS

### ICC as a predictor of species-level social measures

While the ICC was not a strong predictor of group size (Table S1; Fig. 2), the ICC positively predicted the percentage of time that primate species spent grooming. For every one standard deviation increase in the ICC, the odds of grooming increased by 66% (pooled posterior mean = 0.508; 99.2% estimates across iterations > 0; Fig. 2 and Table S2). Models using an ICC calculated from climate data for the exact locations where grooming data was collected, instead of averages over species’ entire geographic ranges, yielded qualitatively similar results (pooled posterior mean = 0.487; 99.2% estimates across iterations > 0; Table S3). Cercopithecidae primates including the Japanese macaque (*Macaca fuscata*), Arunachal macaque (*Macaca munzala*), chacma baboon (*Papio urinus*), Nepal gray langur (*Semnopithecus schistaceus*), and gelada (*Theropithecus gelada*) scored highest on both the percentage of time they spent grooming and the ICC, while a number of langurs and apes like the Mentawai langur (*Presbytis potenziani*), maroon langur (*Presbytis rubicunda*), Kloss’s gibbon (*Hylobates klossii*), Sumatran orangutan (*Pongo abelii*), and Bornean orangutan (*Pongo pygmaeus*) had the lowest combined scores for percent time spent grooming and the ICC (Fig. 3).

**Fig. 2:**
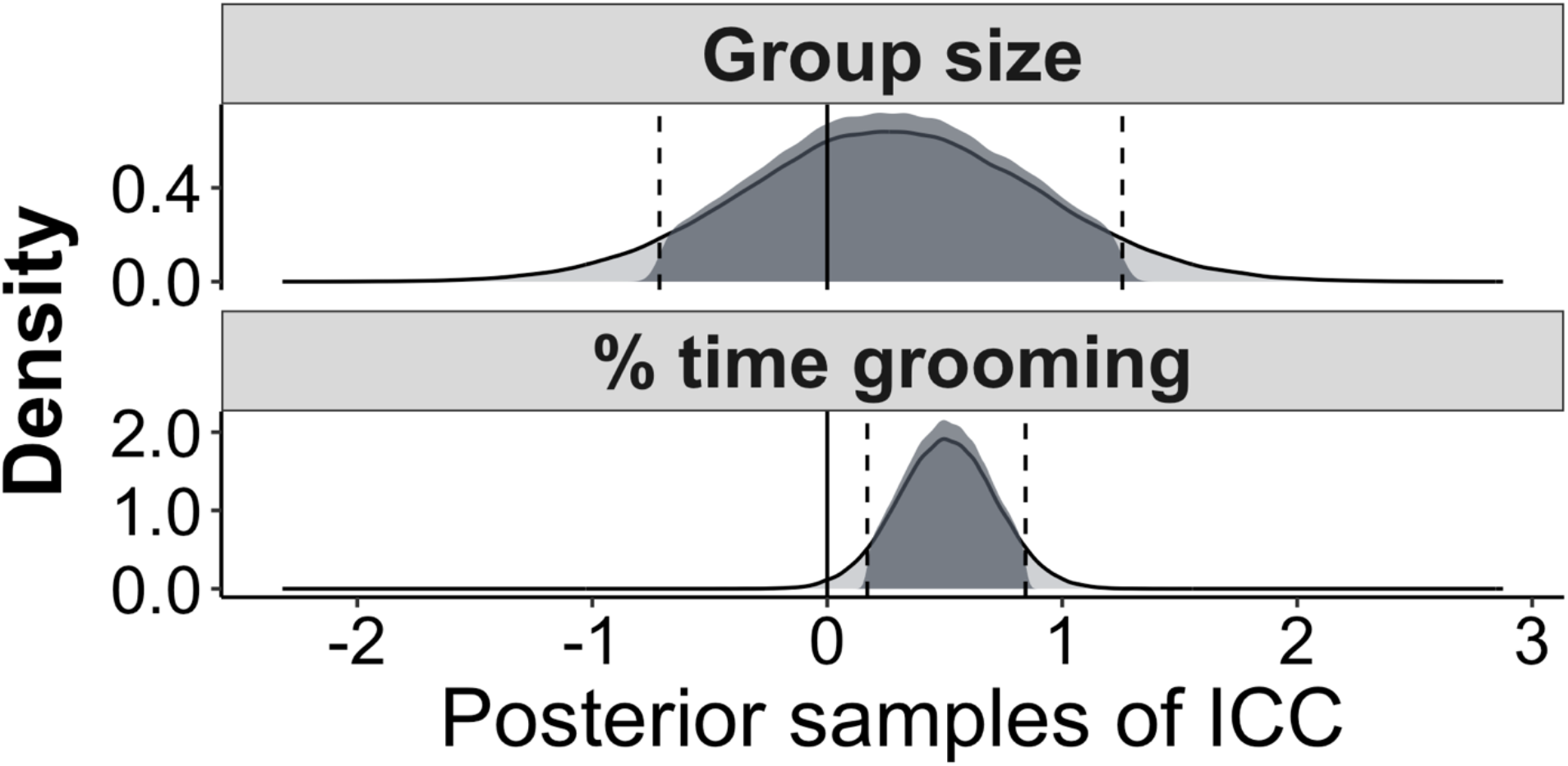
Density of posterior samples of the ICC when modeled as a predictor of two species-level social measures: mean group size and the average percentage of time individuals spend grooming. Each cell contains the density of posterior samples of the ICC for the social outcome across retained iterations from 100 MCMCglmm models run with 100 randomly sampled phylogenies from Upham et al. (2019). The dashed lines and shading represent the 89% credible intervals for these samples.

**Fig. 3:**
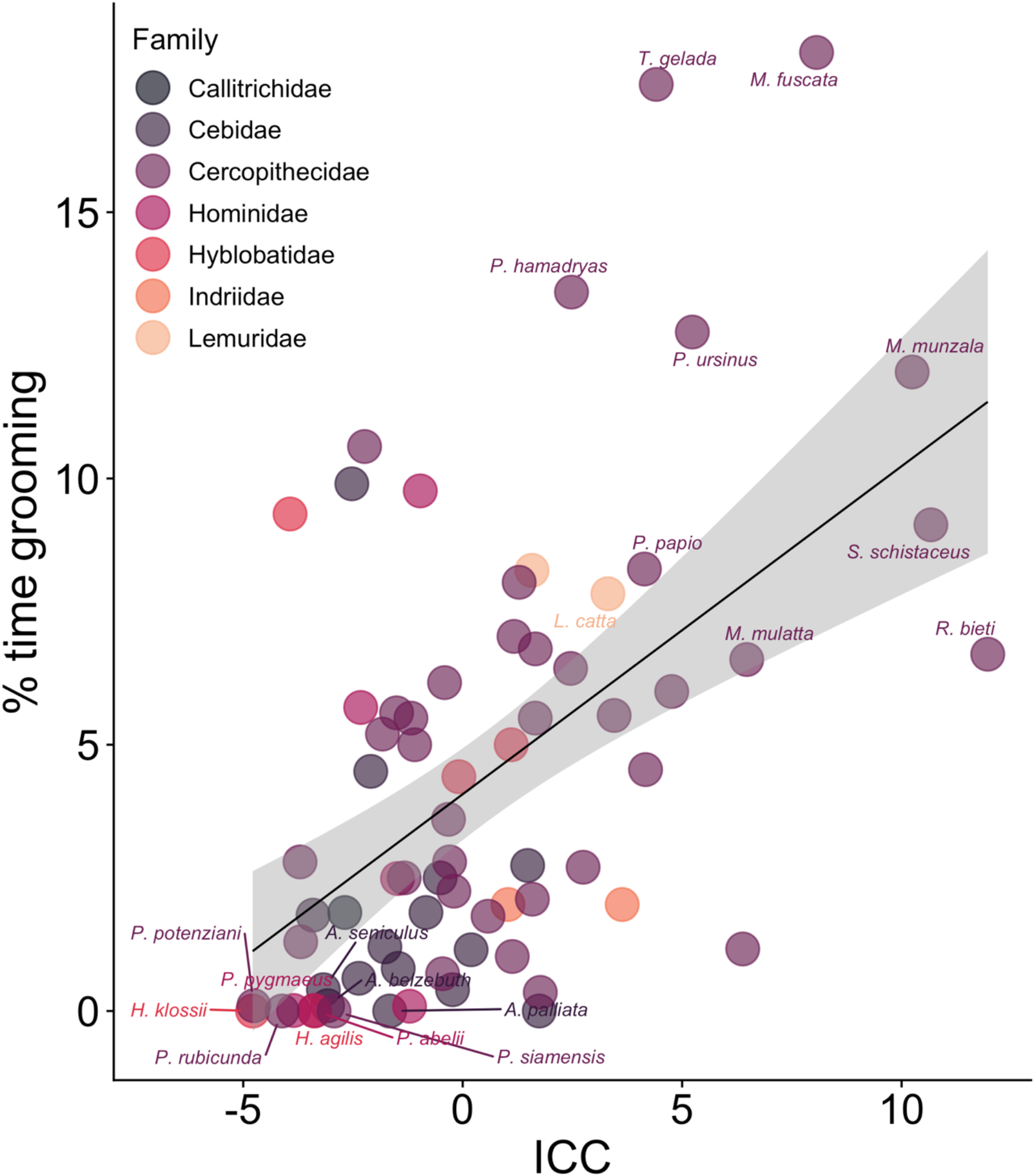
Average percent time individuals spend grooming versus the ICC with colors corresponding to family. Species with the 10 lowest and 10 highest combined scores for both variables are labeled with species names.

When testing the four collinear climate variables used to build the ICC as independent predictors of time spent grooming, reflected mean precipitation was the strongest predictors of time spent grooming (pooled posterior mean = 0.674; 99.9% estimates across iterations > 0; Table S6) followed by temperature seasonality (pooled posterior mean = 0.367; 96.1% estimates across iterations > 0; Table S5). Deviation of mean temperature and precipitation seasonality by contrast had weaker associations (pooled posterior mean = 0.152; 81.0% estimates across iterations > 0; Table S4 and pooled posterior mean = 0.282; 90.9% estimates across iterations > 0; Table S7, respectively). Notably, using mean temperature instead of mean temperature deviation increased the estimated effects of temperature suggesting colder climates in particular had more grooming, but this effect was still smaller than for reflected mean precipitation or temperature seasonality (pooled posterior mean =-0.302; 95.6% estimates across iterations > 0; Table S8). Temperature and precipitation variability across years yielded quantitatively similar results to seasonality within years (Tables S9 and S10; Fig. S3), suggesting short-term and long-term variability in climate have similar effects on the expression of grooming. Consistent with the ICC, no individual climate variables were meaningful predictors of group size (Table S11 to S16).

#### Social organization and parasite analysis

In a model that included group size, female philopatry and social system as predictors, mean group size and pair living both predicted more time spent grooming (pooled posterior mean = 0.699; 99.7% estimates across iterations > 0 and pooled posterior mean = 2.000; 99.2% estimates across iterations > 0, respectively; Table S17). However, even controlling for these variables, percentage time spent grooming and the ICC still shared a strong positive relationship (pooled posterior mean = 0.436; 98.2% estimates across iterations > 0; Table S17), suggesting these aspects of social organization do not mediate the grooming-ICC relationship. When assessing correlations with dominance in a small sample of species, percent time spent grooming and the ICC were both negatively correlated with aggression symmetry (r=-0.33 and r=-0.19, respectively; Fig. S4a and S4b) and counteraggression (r=-0.66 and r=-0.20, respectively, S4c and S4d). These correlations are consistent with the idea that more despotic societies, characterized by lower aggression symmetry and counteraggression, emerge in more challenging climates and favor more time spent grooming, although more data are needed for a formal analysis.

Including the number of ectoparasite types known to affect primate species in models where percent time spent grooming was the response did not meaningfully change the effect of the ICC (pooled posterior mean = 0.472; 98.1% estimates across iterations > 0; Table S18).

## DISCUSSION

Analyses revealed a strong positive association between the average percentage of time individuals spend social grooming and the climatic challenges in species’ environments. In physiologically challenging environments, characterized by low and heterogeneous rainfall and by extreme and heterogenous temperatures, species spent more time socially grooming. Of the four positively correlated climate variables incorporated in this climate index, inverse mean precipitation (i.e., the degree to which an environment is dry) had the largest individual influence on time spent grooming. No effect was found of the same climate measures on average group size suggesting the dimensions of climatic challenges captured by the ICC explain social behavior but not the tendency of animals to live in larger groups.

The mechanisms that could link increased time spent grooming to challenging environments are not mutually exclusive. Here, I discuss in greater detail each of these possibilities, citing existing support for each. First, it is possible that the functional benefits of grooming are greater in challenging environments if, for example, there is increased ectoparasite richness in these species’ environments or grooming helps with thermoregulation. Modeling scenarios suggest that climate change, which is projected to increase climate extremes and decrease predictability, favors a substantial increase in suitable habitats for ticks, one type of ectoparasite, globally (Cumming & Van Vuuren 2006). However, using a limited dataset on ectoparasites available from the GMPD (Nunn & Altizer, 2005), I found no evidence that the number of ectoparasite types for which a species is host explains this relationship. Notably, high ectoparasite abundances are also often recorded in warm, humid, environments (Süss et al. 2008) like rainforests, which tend to score as less challenging on the index used in this analysis. The future development of more comprehensive datasets detailing parasite abundances and primate-parasite interactions would allow for a more robust analysis. In terms of thermoregulation, McFarland et al. (2016) demonstrated that backcombed pelts offer enhanced thermal performance, thus thermoregulation could also play a role in linking grooming and climate. However, total precipitation was a stronger predictor of grooming than temperature, suggesting this is unlikely to be the only factor favoring grooming in challenging climates.

Second, it is possible that the psychological benefits of close affiliative relationships are greater in physiologically challenging climates. Close affiliative relationships, known in primates to be established and maintained via grooming, are demonstrated to buffer against climate induced stress: wild Barbary macaques (*Macaca sylvanus*) with the strongest affiliative relationships have lower fecal glucocorticoid metabolite levels in cold temperatures than those with weaker relationships, even after accounting for the thermoregulatory and immediate functional benefits of social interactions (Young et al. 2014). It is thus possible that affiliative behaviors are expressed more frequently in challenging conditions as individuals seek comfort and support from social partners.

Third, increased competition in challenging environments may favor investment in close social partners. For instance, if challenging climates intensify within-group competition, these conditions may favor investment in coalition partners in the form of grooming (Barton et al. 1996). This theory is supported by the fact that inverse mean annual precipitation—a key determinant of resource availability and distribution—was the climate variable with the largest individual influence on time spent grooming. Moreover, if challenging conditions favor higher risk of infanticide, females may also invest more time in social relationships with males (Henzi & Barrett 2003). In either of these scenarios challenging climates may lead to more time spent grooming overall.

Social organizations and social systems that intensify selection for affiliative behavior could also mediate a relationship between grooming and climatic variables if social organizations or systems that favor more grooming are more common in challenging climates. I explored whether group size, broad classifications of social systems, or female philopatry mediated the grooming-ICC relationship and found no support. However, the role of other aspects of social organization or systems cannot be ruled out. Qi et al. (2023) demonstrated that hierarchal societies are associated with both cold climates and a greater amount of time spent social grooming among Asian colobines. While precipitation was the strongest predictor of social grooming in the present dataset, this association could still be mediated by the emergence of social hierarchies. For instance, more despotic societies may evolve in more challenging climates if these climates have higher feeding competition due to there being fewer and patchier resources, which may lead to steeper social hierarchies and despotic dynamics (Boone 1992). Affiliative interactions may be frequent in these systems as individuals aim to avoid conflict and recruit social support (Seyfarth 1977). Indeed, many species that scored high on both time spent grooming and the ICC tended to have strict dominance hierarchies (e.g., baboons and macaques) while some with very low scores tended to have less strict hierarchies (e.g., howler monkeys and langurs). Moreover, data correlations from a small sample of species suggested that more despotic societies spend more time grooming and live in more challenging climates than less despotic species. Thus, competition (discussed above) and the emergence of despotic hierarchical societies may well play a role in explaining why social grooming is globally favored in challenging climates.

Lastly, time constraints on other aspects of behavior may increase grooming in challenging climates if, for example, challenging conditions induce forced periods of inactivity that favor socializing. In this case, social grooming would not be directly selected for in challenging climates but instead emerge as a by-product of time budget allowances (see, for instance, the comparative analysis of climate as a predictor of colobine activity budgets in Kraus & Strier 2022). Across primate species the average percentage of total time budget observed resting is positively predicted by temperature: individuals of species in hotter climates rest more (Korstjens et al. 2010), as would be expected if challenging conditions favor more time resting. Testing whether time budget trade-offs explain the relationship between grooming and ICC is complicated by the fact that, while foraging time is thought to be the highest priority for wild primates, the causal hierarchical relationship between time resting and grooming is not obvious (Dunbar 1992). Thus, even if time spent resting were correlated with climate variables it would be hard to know whether i) more time resting causes more time grooming, ii) increased grooming leads to more resting, or iii) they are both advantageous and thus both more prevalent in challenging climates.

As part of their investigation of the origins of human social grooming, Jaeggi et al. (2017) analyzed the data in Grueter et al. (2013) using a different framework and found little to no influence of climate/climate-related variables (mean diurnal temperature range, isothermality, annual precipitation, latitude, and altitude) on social grooming. Several differences in data and study design could explain the differences in results reported here. One difference is that Jaeggi et al. (2017) scored terrestriality following Grueter et al. (2013) as opposed to the larger compilation used here, which scored six species differently. However, rescoring these species does not meaningfully influence estimates for the ICC across the models reported above. The use of an older phylogeny and climate data in Jaeggi et al. (2017), or the use of arcsine data transformations and a model selection approach (which can sometimes disadvantage correlated variables) might instead explain the different estimated effects of climate variables in these earlier analyses.

The fact that climate variables did not predict average group size suggests other ecological pressures may be responsible for determining cross-species variation in group size. Obvious examples include variation in the density and distribution of predators (Hamilton, 1971), resource defensibility (e.g., clumped versus dispersed) (Silk, 2007), and/or pathogen transmission (Patterson & Ruckstuhl, 2013). These ecological pressures may be shaped in part by climate, but explain independent variation in group size, and should continue to be explored as determinants of group size in comparative frameworks.

## CONCLUSIONS

Two types of cooperative social behavior—cooperative breeding and social grooming— have now been shown to occur disproportionately in extreme and variable climates. While there are multiple reasons why these behaviors might occur disproportionately under challenging conditions, these results suggest a potential role for climate in influencing the evolution and/or maintenance of cooperative behaviors. Continuing to explore these relationships and their underlying mechanisms in the face of ongoing climate change could lend insights into the future trajectories of social evolution.

## DATA AVAILABILITY STATEMENT

Data and code will be made available upon publication.

## ACKNOWLEDGEMENTS

Thanks to Charles Nunn, Cyril Grueter, Dan Greenberg, Jacob Kraus, Dieter Lukas, Gabriela Venable, Caroline Shearer, Brian Lerch, and the Alberts lab at Duke for their helpful feedback on these results. I also gratefully acknowledge financial support from the Natural Sciences and Engineering Research Council of Canada (PGSD3 - 577867 – 2023) and Duke University.

## Notes

### Competing Interest Statement

The authors have declared no competing interest.

## REFERENCES

Abatzoglou, J. T., Dobrowski, S. Z., Parks, S. A., & Hegewisch, K. C. (2018). TerraClimate, a high-resolution global dataset of monthly climate and climatic water balance from 1958–2015. Sci. Data, 5(1), 170191.

Akinyi, M. Y., Tung, J., Jeneby, M., Patel, N. B., Altmann, J., & Alberts, S. C. (2013). Role of grooming in reducing tick load in wild baboons (Papio cynocephalus). Anim. Behav., 85(3), 559–568.

Andras, P., Lazarus, J., & Roberts, G. (2007). Environmental adversity and uncertainty favour cooperation. BMC Evol. Biol., 7, 1–8.

Barton, R. A., Byrne, R. W., & Whiten, A. (1996). Ecology, feeding competition and social structure in baboons. Behav. Ecol. Sociobiol., 38(5), 321–329.

Boone, J. L. (1992). Competition, conflict, and the development of social hierarchies. Evol. Hum. Behav., 301–337.

Brooke, M. D. L. (1985). The effect of allopreening on tick burdens of molting eudyptid penguins. Auk, 893–895.

Campbell, L. A., Tkaczynski, P. J., Lehmann, J., Mouna, M., & Majolo, B. (2018). Social thermoregulation as a potential mechanism linking sociality and fitness: Barbary macaques with more social partners form larger huddles. Sci. Rep., 8(1), 6074.

Campos, F. A., Kalbitzer, U., Melin, A. D., Hogan, J. D., Cheves, S. E., Murillo-Chacon, E., et al. (2020). Differential impact of severe drought on infant mortality in two sympatric neotropical primates. R. Soc. Open Sci., 7(4), 200302.

Carrera, S. C., Godoy, I., Gault, C. M., Mensing, A., Damm, J., Perry, S. E. et al. (2025). Stress responsiveness in a wild primate predicts survival across an extreme El Niño drought. Sci. Adv., 11(4), eadq5020.

Cornwallis, C. K., Botero, C. A., Rubenstein, D. R., Downing, P. A., West, S. A., & Griffin, A. S. (2017). Cooperation facilitates the colonization of harsh environments. Nat. Ecol. Evol., 1(3), 0057.

Creel, S., & Creel, N. M. (1995). Communal hunting and pack size in African wild dogs, Lycaon pictus. Anim. Behav., 50(5), 1325–1339.

Creighton, M. J. A., & Nunn, C. L. (2023). Explaining the primate extinction crisis: predictors of extinction risk and active threats. Proc. R. Soc. B, 290(2006), 20231441.

Cumming, G. S., & Van Vuuren, D. P. (2006). Will climate change affect ectoparasite species ranges? Glob. Ecol. Biogeogr., 15(5), 486–497.

Dunbar, R. I. M. (1991). Functional significance of social grooming in primates. Folia Primatol., 57(3), 121–131.

Dunbar, R. I. M. (1992). Time: a hidden constraint on the behavioural ecology of baboons. Behav. Ecol. Sociobiol., 31(1), 35–49.

Dunbar, R. I. M. (1988). Primate social systems. Cornell University Press, Ithaca, New York, USA.

Dunbar, R. I. M., & Lehmann, J. (2013). Grooming and social cohesion in primates: a comment on Grueter et al. Evol. Hum. Behav., 34(6), 453–455.

Estrada, A., Garber, P. A., Gouveia, S., Fernández-Llamazares, Á., Ascensão, F., Fuentes, A., et al. (2022). Global importance of Indigenous Peoples, their lands, and knowledge systems for saving the world’s primates from extinction. Sci. Adv., 8(31), eabn2927.

Fick, S. E., & Hijmans, R. J. (2017). WorldClim 2: new 1-km spatial resolution climate surfaces for global land areas. Int. J. Climatol., 37(12), 4302–4315.

Firman, R. C., Rubenstein, D. R., Moran, J. M., Rowe, K. C., & Buzatto, B. A. (2020). Extreme and variable climatic conditions drive the evolution of sociality in Australian rodents. Curr. Biol., 30(4), 691–697.

Galán-Acedo, C., Arroyo-Rodríguez, V., Andresen, E., & Arasa-Gisbert, R. (2019). Ecological traits of the world’s primates. Sci. Data, 6(1), 55.

Grueter, C. C., Bissonnette, A., Isler, K., & van Schaik, C. P. (2013). Grooming and group cohesion in primates: implications for the evolution of language. Evol. Hum. Behav., 34(1), 61–68.

Hadfield, J. D. (2024). Package ‘MCMCglmm’ (Version 2.36). R package. Available at: https://cran.r-project.org/package=MCMCglmm.

Hamilton, W. D. (1971). Geometry for the selfish herd. J. Theor. Biol., 31(2), 295–311.

Hamilton III, W. J., Buskirk, R. E., & Buskirk, W. H. (1975). Chacma baboon tactics during intertroop encounters. J. Mammal., 56(4), 857–870.

Henzi, P., & Barrett, L. (2003). Evolutionary ecology, sexual conflict, and behavioral differentiation among baboon populations. Evol. Anthropol., 12(5), 217–230.

Hijmans, R. J., Barbosa, M., Ghosh, A., & Mandel, A. (2024). Package ‘geodata’ (Version 0. 6-9). Available at: https://cran.r-project.org/package=geodata.

Jaeggi, A. V., Kramer, K. L., Hames, R., Kiely, E. J., Gomes, C., Kaplan, H., et al. (2017). Human grooming in comparative perspective: People in six small-scale societies groom less but socialize just as much as expected for a typical primate. Am. J. Phys. Anthropol., 162(4), 810–816.

IUCN (2025). The IUCN Red List of Threatened Species (Version 2025-2). Available at: https://www.iucnredlist.org. Last accessed 02 February 2026.

Jetz, W., & Rubenstein, D. R. (2011). Environmental uncertainty and the global biogeography of cooperative breeding in birds. Curr. Biol., 21(1), 72–78.

Kamilar, J. M., & Baden, A. L. (2014). What drives flexibility in primate social organization? Behav. Ecol. Sociobiol., 68(10), 1677–1692.

Kavanagh, E., Street, S. E., Angwela, F. O., Bergman, T. J., Blaszczyk, M. B., Bolt, L. M., et al. (2021). Dominance style is a key predictor of vocal use and evolution across nonhuman primates. R. Soc. Open Sci., 8(7), 210873.

Komdeur, J., & Ma, L. (2021). Keeping up with environmental change: The importance of sociality. Ethology, 127(10), 790–807.

Korstjens, A. H., Lehmann, J., & Dunbar, R. I. M. (2010). Resting time as an ecological constraint on primate biogeography. Anim. Behav., 79(2), 361–374.

Kraus, J. B., & Strier, K. B. (2022). Geographic, climatic, and phylogenetic drivers of variation in colobine activity budgets. Primates, 63(6), 647–658.

Lehmann, J., Korstjens, A. H., & Dunbar, R. I. (2007). Group size, grooming and social cohesion in primates. Anim. Behav., 74(6), 1617–1629.

Lucore, J. M., Beehner, J. C., White, A. F., Sinclair, L. F., Martins, V. A., Kovalaskas, S. A., et al. (2024). High temperatures are associated with decreased immune system performance in a wild primate. Sci. Adv., 10(48), eadq6629.

Lukas, D., & Clutton-Brock, T. (2017). Climate and the distribution of cooperative breeding in mammals. R. Soc. Open Sci., 4(1), 160897.

Majolo, B., McFarland, R., Young, C., & Qarro, M. (2013). The effect of climatic factors on the activity budgets of Barbary macaques (Macaca sylvanus). Int. J. Primatol., 34(3), 500–514.

McFarland, R., & Majolo, B. (2013). Coping with the cold: predictors of survival in wild Barbary macaques, Macaca sylvanus. Biol. Lett., 9(4), 20130428.

McFarland, R., Henzi, S. P., Barrett, L., Wanigaratne, A., Coetzee, E., Fuller, A., … & Maloney, S. K. (2016). Thermal consequences of increased pelt loft infer an additional utilitarian function for grooming. Am. J. Primatol., 78(4), 456–461.

Nunn, C. L., & Altizer, S. M. (2005). The global mammal parasite database: an online resource for infectious disease records in wild primates. Evol. Anthropol., 14(1), 1–2.

Packer, C., Scheel, D., & Pusey, A. E. (1990). Why lions form groups: food is not enough. The Am. Nat., 136(1), 1–19.

Patterson, J. E., & Ruckstuhl, K. E. (2013). Parasite infection and host group size: a meta-analytical review. Parasitology, 140(7), 803–813.

Peters, R. H. (1986). The ecological implications of body size. Vol. 2. Cambridge University Press, Cambridge, Cambridgeshire, England.

Qi, X. G., Wu, J., Zhao, L., Wang, L., Guang, X., Garber, P. A., et al. (2023). Adaptations to a cold climate promoted social evolution in Asian colobine primates. Science, 380(6648), eabl8621.

R Core Team (2025). R: A language and environment for statistical computing (Version 4.5.1, “Great Square Root”). R Foundation for Statistical Computing, Vienna, Austria. Available at: https://www.R-project.org/.

Rowe N., & Myers M. (2011). All the world’s primates. Primate Conservation, Inc. Available at: https://www.alltheworldsprimates.org. Last accessed 16 December 2022.

Russo, D., Cistrone, L., Budinski, I., Console, G., Della Corte, M., Milighetti, C., et al. (2017). Sociality influences thermoregulation and roost switching in a forest bat using ephemeral roosts. Ecol. Evol., 7(14), 5310–5321.

Rubenstein, D. R., & Lovette, I. J. (2007). Temporal environmental variability drives the evolution of cooperative breeding in birds. Curr. Biol., 17(16), 1414–1419.

Seyfarth, R. M. (1977). A model of social grooming among adult female monkeys. J. Theor. Biol., 65(4), 671–698.

Silk, J. B. (2007). The adaptive value of sociality in mammalian groups. Philos. Trans. R. Soc. B., 362(1480), 539–559.

Smaldino, P. E., Schank, J. C., & McElreath, R. (2013). Increased costs of cooperation help cooperators in the long run. Am. Nat., 181, 451–463.

Srivastava, D. S., & Lawton, J. H. (1998). Why more productive sites have more species: an experimental test of theory using tree-hole communities. Am. Nat., 152(4), 510–529.

Süss, J., Klaus, C., Gerstengarbe, F. W., & Werner, P. C. (2008). What makes ticks tick? Climate change, ticks, and tick-borne diseases. J. Travel Med., 15(1), 39–45.

Tattersall, G. J., Sinclair, B. J., Withers, P. C., Fields, P. A., Seebacher, F., Cooper, C. E., et al. (2012). Coping with thermal challenges: physiological adaptations to environmental temperatures. Compr. Physiol., 2(3), 2151–2202.

Taylor, S. E., Klein, L. C., Gruenewald, T. L., Gurung, R. A., & Fernandes-Taylor, S. (2003). Affiliation, social support, and biobehavioral responses to stress. In: Social psychological foundations of health and illness, eds. Suls, J. & Wallston, K. A. Blackwell Publishing, Malden, USA, pp. 314–331.

Upham, N. S., Esselstyn, J. A., & Jetz, W. (2019). Inferring the mammal tree: species-level sets of phylogenies for questions in ecology, evolution, and conservation. PLoS Biol., 17(12), e3000494.

van Schaik, C. P., & Pfannes, K. R. (2005). Tropical climates and phenology. In: Seasonality in primates: Studies of living and extinct human and non-human primates, eds. Brockman, D. K. & van Schaik, C. P. Cambridge University Press, Cambridge, Cambridgeshire, England, pp. 23–54.

Villa, S. M., Goodman, G. B., Ruff, J. S., & Clayton, D. H. (2016). Does allopreening control avian ectoparasites? Biol. Lett., 12(7), 20160362.

White, T. (2008). The role of food, weather and climate in limiting the abundance of animals. Biol. Rev., 83(3), 227–248.

Wittig, R. M., Crockford, C., Lehmann, J., Whitten, P. L., Seyfarth, R. M., & Cheney, D. L. (2008). Focused grooming networks and stress alleviation in wild female baboons. Horm. Behav., 54(1), 170–177.

Young, C., Majolo, B., Heistermann, M., Schülke, O., & Ostner, J. (2014). Responses to social and environmental stress are attenuated by strong male bonds in wild macaques. Proc. Natl. Acad. Sci. U. S. A., 111(51), 18195–18200.

Young, C., Bonnell, T. R., Brown, L. R., Dostie, M. J., Ganswindt, A., Kienzle, S., et al. (2019). Climate induced stress and mortality in vervet monkeys. R. Soc. Open Sci., 6(11), 191078.

