## Supplementary Materials for "Species-level variation in primate social behavior is correlated with climate extremes and variability"

### SUPPLEMENTARY METHODS

*Phylogenetic signal.* The mean phylogenetic signal of both social traits of interest (average group size and percent time spent grooming) was computed as Pagel’s Lambda (λ) (Pagel, 1999) using the phylosig function from ‘phytools’ (Revell, 2024), using the random sample of 100 trees sourced from Upham et al. (2019) to account for phylogenetic uncertainty. A λ of zero suggests a trait is randomly distributed with respect to phylogeny while a λ closer to one suggests greater phylogenetic dependence.

*Social organization and system.* Data for several aspects of social organization/system that could mediate a relationship between percent time spent grooming and the ICC were sourced from existing compilations. Data on social system (solitary, pair living, polygyny, and polygynandry) were taken for species from the compilation in Creighton & Nunn (2023). Social system data in this compilation came primarily from DeCasien et al. (2017) and were supplemented with information from Rowe & Myers (2011) and other sources. Notably, while these categories are typically used to refer to mating system, they are used in DeCasien et al. (2017) to describe social systems in addition to mating systems.

Data on female philopatry (true=1 and false=0) were obtained for all 70 species with grooming data from Lehmann et al. (2007) when available, and supplemented or updated using accounts from other sources. Species were scored as female philopatric if they exhibited sex-biased female philopatry without significant female movement. For species where dispersal patterns were unknown, philopatry was inferred based on dispersal patterns of close relatives in the same genus.

These data should be updated as more information on dispersal becomes available for lesser studied populations.

Data on aggression symmetry and counteraggression were taken from Kavanagh et al. (2021). This compilation reports dominance styles for 26 primate species. Aggression symmetry was measured using the directional inconsistency index, where values reflect the proportion of aggression bouts across dyads where the role of the aggressor and victim occurred in the least frequent direction (Kavanagh et al. 2021). Counteraggression was measured as the percent of aggressive bouts that were followed by retaliation from the victim (Kavanagh et al. 2021). More despotic species tend to have lower aggression symmetry and less counteraggression, reflecting the imbalance in social capita among individuals. Of the 70 species with social grooming time data, eleven in Kavanagh et al. (2021) had data on aggression symmetry and eight had data on rates of counteraggression.

*Parasite counts*. To test if a greater ectoparasite burden explains the relationship between the ICC and grooming, one would ideally obtain estimates of the total density of ectoparasites in each species' habitat for which the species is an eligible host. However, these data do not exist at the species-level or for enough primate species in the published literature to create a compilation. Thus, for the purposes of conducting an exploratory analysis I resorted to using the most complete and taxonomically comprehensive compilation available, the Global Mammal Parasite Database (GMPD) (Nunn & Altizer, 2005), which provides information about the number of different ectoparasite types that are known to affect each primate species. From this database I retrieved a count of the number of parasites of the type "arthropod" (i.e., ectoparasites) known to affect each primate species. Sampling effort (i.e., the number of papers published on each species recorded in the GMPD) was taken from Herrera et al. (2023) to account for biases in the number of parasites reported to affect each species caused by variation in research effort. 81 primate species in my dataset were documented to be affected by between one and 15 ectoparasite types. Species that had no recorded ectoparasites but did have sampling effort in the GMPD were scored as have zero ectoparasites in my dataset while species in the dataset with no research effort in the GMPD were scored as missing data. Of the 70 species in my dataset with grooming data, 55 had non-missing ectoparasite data. Further commentary on the biases and limitations associated with these parasite data for inferring differences in ectoparasite burdens across species is presented in the main text.

### SUPPLEMENTARY RESULTS

*Phylogenetic signal.* Average group size and percent time spent grooming were both phylogenetically clustered. Across 100 phylogenies the mean λ for group size was 0.752 (95% CI = 0.629 to 0.995) and the mean λ for percent time spent grooming was 0.806 (95% CI = 0.740 to 0.858), meaning relatively high phylogenetic signal.

### SUPPLEMENTARY TABLES

**Table S1:** Summarized results from 100 MCMCglmm models with a Gaussian distribution predicting average group size across 259 primate species where the Index of Climatic Challenges (ICC), average adult body mass, and terrestrial (true or false) were included as fixed effects. All continuous fixed effects were centred with respect to their means and scaled by one standard deviation.

| **term** | **mean posterior mean** | **mean lower 89% CI** | **mean upper 89% CI** | **%coef +** | **%coef -** |
|---|---|---|---|---|---|
| Intercept | -0.381 | -2.714 | 1.914 | 39.7 | 60.3 |
| ICC | 0.270 | -0.710 | 1.259 | 66.8 | 33.2 |
| Body mass | 0.450 | -0.611 | 1.493 | 75.3 | 24.7 |
| Terrestrial | 2.056 | -0.302 | 4.436 | 91.7 | 8.3 |

**Table S2:** Summarized results from 100 MCMCglmm models with a Gaussian distribution predicting logit-transformed average percent time spent grooming across 70 primate species where the ICC, average adult body mass, and terrestrial (true or false) were included as fixed effects. All continuous fixed effects were centred with respect to their means and scaled by one standard deviation.

| term | mean posterior mean | mean lower 89% CI | mean upper 89% CI | %coef + | %coef - |
|---|---|---|---|---|---|
| Intercept | -2.127 | -4.079 | -0.270 | 4.4 | 95.6 |
| ICC | 0.508 | 0.177 | 0.850 | 99.2 | 0.8 |
| Body mass | -0.380 | -0.801 | 0.029 | 7.3 | 92.7 |
| Terrestrial | 0.546 | -0.197 | 1.304 | 87.9 | 12.1 |

**Table S3:** Summarized results from 100 MCMCglmm models with a Gaussian distribution predicting logit-transformed percent time spent grooming across 70 primate species where a field site-specific ICC, average adult body mass, and terrestrial (true or false) were included as fixed effects. All continuous fixed effects were centred with respect to their means and scaled by one standard deviation.

| term | mean posterior mean | mean lower 89% CI | mean upper 89% CI | %coef + | %coef - |
|---|---|---|---|---|---|
| Intercept | -2.465 | -4.374 | -0.639 | 2.6 | 97.4 |
| Site-specific ICC | 0.487 | 0.165 | 0.809 | 99.2 | 0.8 |
| Body mass | -0.446 | -0.849 | -0.047 | 4.2 | 95.8 |
| Terrestrial | 0.757 | 0.042 | 1.458 | 95.6 | 4.4 |

**Table S4:** Summarized results from 100 MCMCglmm models with a Gaussian distribution predicting logit-transformed average percent time spent grooming across 70 primate species where deviation of mean temperature, average adult body mass, and terrestrial (true or false) were included as fixed effects. All continuous fixed effects were centred with respect to their means and scaled by one standard deviation.

| **term** | **mean posterior mean** | **mean lower 89% CI** | **mean upper 89% CI** | **%coef +** | **%coef -** |
|---|---|---|---|---|---|
| Intercept | -2.138 | -4.097 | -0.282 | 4.2 | 95.8 |
| Deviation of mean temperature | 0.152 | -0.126 | 0.432 | 81.0 | 19.0 |
| Body mass | -0.397 | -0.835 | 0.032 | 7.4 | 92.6 |
| Terrestrial | 0.918 | 0.163 | 1.649 | 97.5 | 2.5 |

**Table S5:** Summarized results from 100 MCMCglmm models with a Gaussian distribution predicting logit-transformed percent time spent grooming across 70 primate species where temperature seasonality, average adult body mass, and terrestrial (true or false) were included as fixed effects. All continuous fixed effects were centred with respect to their means and scaled by one standard deviation.

| **term** | **mean posterior mean** | **mean lower 89% CI** | **mean upper 89% CI** | **%coef +** | **%coef -** |
|---|---|---|---|---|---|
| Intercept | -2.030 | -3.957 | -0.160 | 4.9 | 95.1 |
| Temperature seasonality | 0.367 | 0.034 | 0.698 | 96.1 | 3.9 |
| Body mass | -0.368 | -0.808 | 0.062 | 9.0 | 91.0 |
| Terrestrial | 0.762 | 0.035 | 1.500 | 95.2 | 4.8 |

**Table S6:** Summarized results from 100 MCMCglmm models with a Gaussian distribution predicting logit-transformed average percent time spent grooming across 70 primate species where reflected mean precipitation, average adult body mass, and terrestrial (true or false) were included as fixed effects. All continuous fixed effects were centred with respect to their means and scaled by one standard deviation.

| **term** | **mean posterior mean** | **mean lower 89% CI** | **mean upper 89% CI** | **%coef +** | **%coef -** |
|---|---|---|---|---|---|
| Intercept | -2.408 | -4.380 | -0.593 | 3.2 | 96.8 |
| Reflected mean precipitation | 0.674 | 0.350 | 1.002 | 99.9 | 0.1 |
| Body mass | -0.418 | -0.814 | -0.026 | 5.0 | 95.0 |
| Terrestrial | 0.443 | -0.296 | 1.174 | 83.2 | 16.8 |

**Table S7:** Summarized results from 100 MCMCglmm models with a Gaussian distribution predicting logit-transformed average percent time spent grooming across 70 primate species where precipitation seasonality, average adult body mass, and terrestrial (true or false) were included as fixed effects. All continuous fixed effects were centred with respect to their means and scaled by one standard deviation.

| **term** | **mean posterior mean** | **mean lower 89% CI** | **mean upper 89% CI** | **%coef +** | **%coef -** |
|---|---|---|---|---|---|
| Intercept | -2.361 | -4.367 | -0.533 | 3.2 | 96.8 |
| Precipitation seasonality | 0.282 | -0.058 | 0.624 | 90.9 | 9.1 |
| Body mass | -0.375 | -0.795 | 0.045 | 7.9 | 92.1 |
| Terrestrial | 0.827 | 0.076 | 1.590 | 95.9 | 4.1 |

**Table S8:** Summarized results from 100 MCMCglmm models with a Gaussian distribution predicting logit-transformed average percent time spent grooming across 70 primate species where mean temperature (rather than deviation in mean temperature), average adult body mass, and terrestrial (true or false) were included as fixed effects. All continuous fixed effects were centred with respect to their means and scaled by one standard deviation.

| term | mean posterior mean | mean lower 89% CI | mean upper 89% CI | %coef + | %coef - |
|---|---|---|---|---|---|
| Intercept | -2.012 | -3.951 | -0.144 | 5.1 | 94.9 |
| Mean temperature | -0.302 | -0.581 | -0.018 | 4.4 | 95.6 |
| Body mass | -0.416 | -0.855 | 0.017 | 6.6 | 93.4 |
| Terrestrial | 0.830 | 0.118 | 1.563 | 96.7 | 3.3 |

**Table S9:** Summarized results from 100 MCMCglmm models with a Gaussian distribution predicting logit-transformed average percent time spent grooming across 70 primate species where across-year temperature variability, average adult body mass, and terrestrial (true or false) were included as fixed effects. All continuous fixed effects were centred with respect to their means and scaled by one standard deviation.

| term | mean posterior mean | mean lower 89% CI | mean upper 89% CI | %coef + | %coef - |
|---|---|---|---|---|---|
| Intercept | -2.171 | -4.084 | -0.361 | 3.8 | 96.2 |
| Across-year temperature variability | 0.383 | 0.109 | 0.653 | 98.8 | 1.2 |
| Body mass | -0.422 | -0.844 | -0.001 | 5.8 | 94.2 |
| Terrestrial | 0.781 | 0.086 | 1.502 | 96.1 | 3.9 |

**Table S10:** Summarized results from 100 MCMCglmm models with a Gaussian distribution predicting logit-transformed average percent time spent grooming across 70 primate species where across-year precipitation variability, average adult body mass, and terrestrial (true or false) were included as fixed effects. All continuous fixed effects were centred with respect to their means and scaled by one standard deviation.

| term | mean posterior mean | mean lower 89% CI | mean upper 89% CI | %coef + | %coef - |
|---|---|---|---|---|---|
| Intercept | -2.184 | -4.116 | -0.289 | 4.0 | 96.0 |
| Across-year precipitation variability | -0.002 | -0.294 | 0.291 | 49.7 | 50.3 |
| Body mass | -0.392 | -0.838 | 0.045 | 8.1 | 91.9 |
| Terrestrial | 1.002 | 0.241 | 1.759 | 98.1 | 1.9 |

**Table S11:** Summarized results from 100 MCMCglmm models with a Gaussian distribution predicting average group size across 259 primate species where deviation of mean temperature, average adult body mass, and terrestrial (true or false) were included as fixed effects. All continuous fixed effects were centred with respect to their means and scaled by one standard deviation.

| term | mean posterior mean | mean lower 89% CI | mean upper 89% CI | %coef + | %coef - |
|---|---|---|---|---|---|
| Intercept | -0.392 | -2.729 | 1.897 | 39.2 | 60.8 |
| Deviation of mean temperature | -0.172 | -1.105 | 0.793 | 38.5 | 61.5 |
| Body mass | 0.458 | -0.576 | 1.534 | 75.6 | 24.4 |
| Terrestrial | 2.095 | -0.218 | 4.509 | 92.1 | 7.9 |

**Table S12:** Summarized results from 100 MCMCglmm models with a Gaussian distribution predicting average group size across 259 primate species where temperature seasonality, average adult body mass, and terrestrial (true or false) were included as fixed effects. All continuous fixed effects were centred with respect to their means and scaled by one standard deviation.

| term | mean posterior mean | mean lower 89% CI | mean upper 89% CI | %coef + | %coef - |
|---|---|---|---|---|---|
| Intercept | -0.391 | -2.698 | 1.913 | 39.4 | 60.6 |
| Temperature seasonality | 0.131 | -0.865 | 1.123 | 58.3 | 41.7 |
| Body mass | 0.454 | -0.605 | 1.502 | 75.7 | 24.3 |
| Terrestrial | 2.065 | -0.300 | 4.437 | 91.8 | 8.2 |

**Table S13:** Summarized results from 100 MCMCglmm models with a Gaussian distribution predicting average group size across 259 primate species where reflected mean precipitation, average adult body mass, and terrestrial (true or false) were included as fixed effects. All continuous fixed effects were centred with respect to their means and scaled by one standard deviation.

| term | mean posterior mean | mean lower 89% CI | mean upper 89% CI | %coef + | %coef - |
|---|---|---|---|---|---|
| Intercept | -0.380 | -2.672 | 1.962 | 39.8 | 60.2 |
| Reflected mean precipitation | 0.356 | -0.625 | 1.339 | 71.8 | 28.2 |
| Body mass | 0.452 | -0.601 | 1.499 | 75.5 | 24.5 |
| Terrestrial | 2.038 | -0.343 | 4.382 | 91.6 | 8.4 |

**Table S14:** Summarized results from 100 MCMCglmm models with a Gaussian distribution predicting average group size across 259 primate species where precipitation seasonality, average adult body mass, and terrestrial (true or false) were included as fixed effects. All continuous fixed effects were centred with respect to their means and scaled by one standard deviation.

| term | mean posterior mean | mean lower 89% CI | mean upper 89% CI | %coef + | %coef - |
|---|---|---|---|---|---|
| Intercept | -0.381 | -2.709 | 1.916 | 39.6 | 60.4 |
| Precipitation seasonality | 0.497 | -0.490 | 1.522 | 78.4 | 21.6 |
| Body mass | 0.460 | -0.601 | 1.496 | 75.8 | 24.2 |
| Terrestrial | 2.049 | -0.350 | 4.370 | 91.7 | 8.3 |

**Table S15:** Summarized results from 100 MCMCglmm models with a Gaussian distribution predicting average group size across 259 primate species where across-year temperature variability, average adult body mass, and terrestrial (true or false) were included as fixed effects. All continuous fixed effects were centred with respect to their means and scaled by one standard deviation.

| term | mean posterior mean | mean lower 89% CI | mean upper 89% CI | %coef + | %coef - |
|---|---|---|---|---|---|
| Intercept | -0.395 | -2.720 | 1.899 | 39.3 | 60.7 |
| Across-year temperature variability | 0.115 | -0.837 | 1.072 | 57.6 | 42.4 |
| Body mass | 0.454 | -0.579 | 1.518 | 75.5 | 24.5 |
| Terrestrial | 2.078 | -0.274 | 4.458 | 91.9 | 8.1 |

**Table S16:** Summarized results from 100 MCMCglmm models with a Gaussian distribution predicting average group size across 259 primate species where across-year precipitation variability, average adult body mass, and terrestrial (true or false) were included as fixed effects. All continuous fixed effects were centred with respect to their means and scaled by one standard deviation.

| **term** | **mean posterior mean** | **mean lower 89% CI** | **mean upper 89% CI** | **%coef +** | **%coef -** |
|---|---|---|---|---|---|
| Intercept | -0.390 | -2.709 | 1.909 | 39.6 | 60.4 |
| Across-year precipitation variability | -0.025 | -0.991 | 0.957 | 48.4 | 51.6 |
| Body mass | 0.453 | -0.595 | 1.503 | 75.6 | 24.4 |
| Terrestrial | 2.090 | -0.244 | 4.480 | 92.1 | 7.9 |

**Table S17:** Summarized results from 100 MCMCglmm models with a Gaussian distribution predicting logit-transformed average percent time spent grooming across 62 primate species where the ICC, average adult body mass, and terrestrial (true or false), mean group size, social system (baseline: Solitary), and female philopatry (true or false) were included as fixed effects. All continuous fixed effects were centred with respect to their means and scaled by one standard deviation.

| **term** | **mean posterior mean** | **mean lower 89% CI** | **mean upper 89% CI** | **%coef +** | **%coef -** |
|---|---|---|---|---|---|
| Intercept | -4.367 | -6.183 | -2.668 | 0.1 | 99.9 |
| ICC | 0.436 | 0.115 | 0.770 | 98.2 | 1.8 |
| Body mass | -0.136 | -0.476 | 0.214 | 26.3 | 73.7 |
| Terrestrial | -0.061 | -0.796 | 0.628 | 44.4 | 55.6 |
| Mean group size | 0.699 | 0.306 | 1.088 | 99.7 | 0.3 |
| Social system: Pair living | 2.000 | 0.726 | 3.273 | 99.2 | 0.8 |
| Social system: Polygyny | 0.786 | -0.410 | 1.970 | 85.4 | 14.6 |
| Social system: Polygynandry | 1.252 | -0.005 | 2.464 | 94.5 | 5.5 |
| Female philopatry | 0.228 | -0.370 | 0.813 | 73.4 | 26.6 |

**Table S18:** Summarized results from 100 MCMCglmm models with a Gaussian distribution predicting logit-transformed average percent time spent grooming across 55 primate species where the ICC, average adult body mass, and terrestrial (true or false), ectoparasite types, and sampling effort were included as fixed effects. All continuous fixed effects were centred with respect to their means and scaled by one standard deviation.

| **term** | **mean posterior mean** | **mean lower 89% CI** | **mean upper 89% CI** | **%coef +** | **%coef -** |
|---|---|---|---|---|---|
| Intercept | -2.249 | -4.205 | -0.410 | 4.0 | 96.0 |
| ICC | 0.472 | 0.109 | 0.834 | 98.1 | 1.9 |
| Body mass | -0.536 | -1.004 | -0.080 | 3.7 | 96.3 |
| Terrestrial | 0.519 | -0.284 | 1.329 | 84.9 | 15.1 |
| Ectoparasite types | -0.184 | -0.557 | 0.183 | 21.2 | 78.8 |
| Sampling effort | 0.297 | -0.099 | 0.681 | 89.0 | 11.0 |

### SUPPLEMENTARY FIGURES

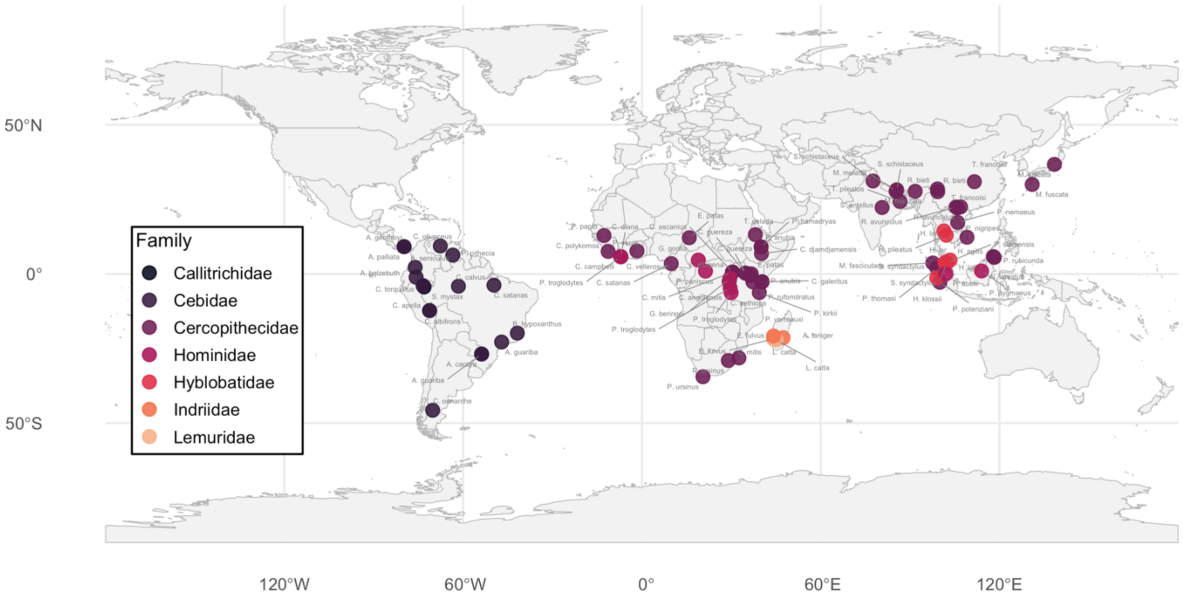


**Fig. S1:** Site coordinates for study populations across 70 species in the dataset with percent time spent grooming data sourced from Grueter et al. (2013). Each species is represented by between one and three study populations.

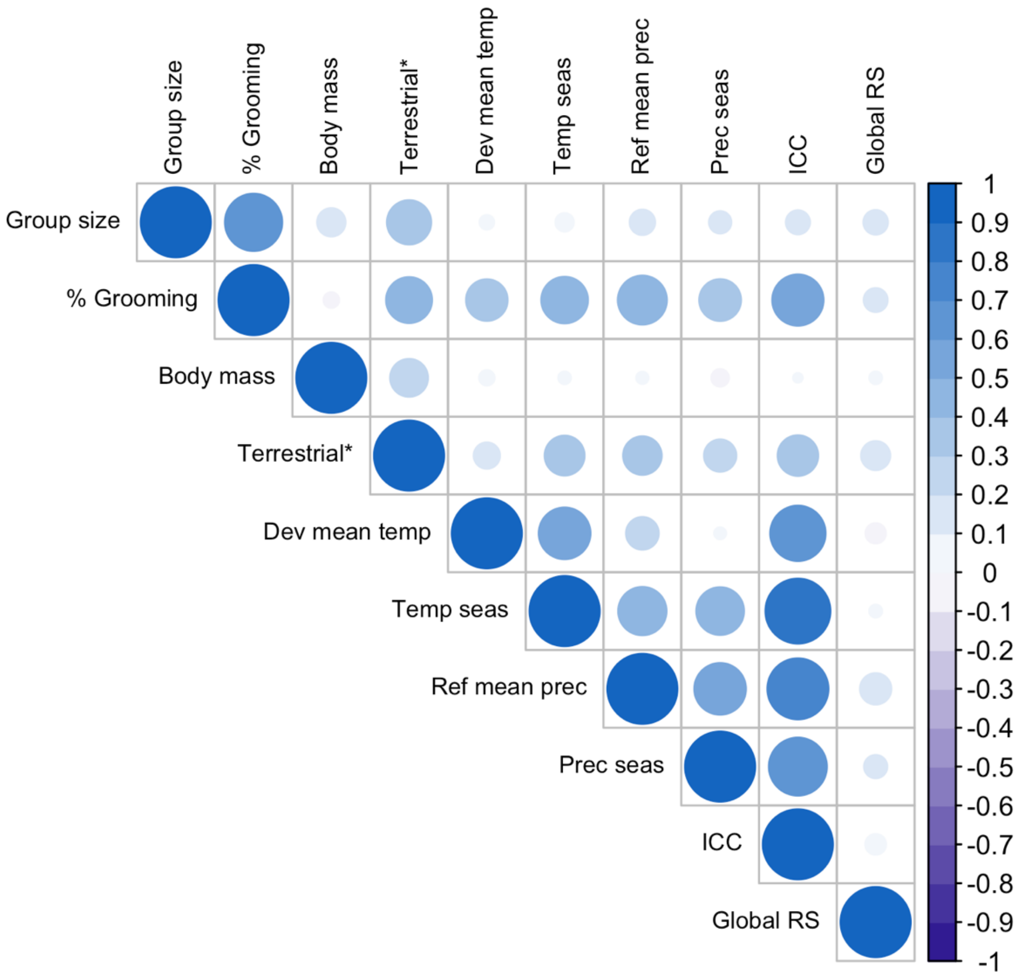


**Fig. S2:** Correlation matrix of species-level social measure, climate variables from species' geographic ranges, and global range size of species' known habitat. Variables indicated with a * were binary variables as opposed to continuous.

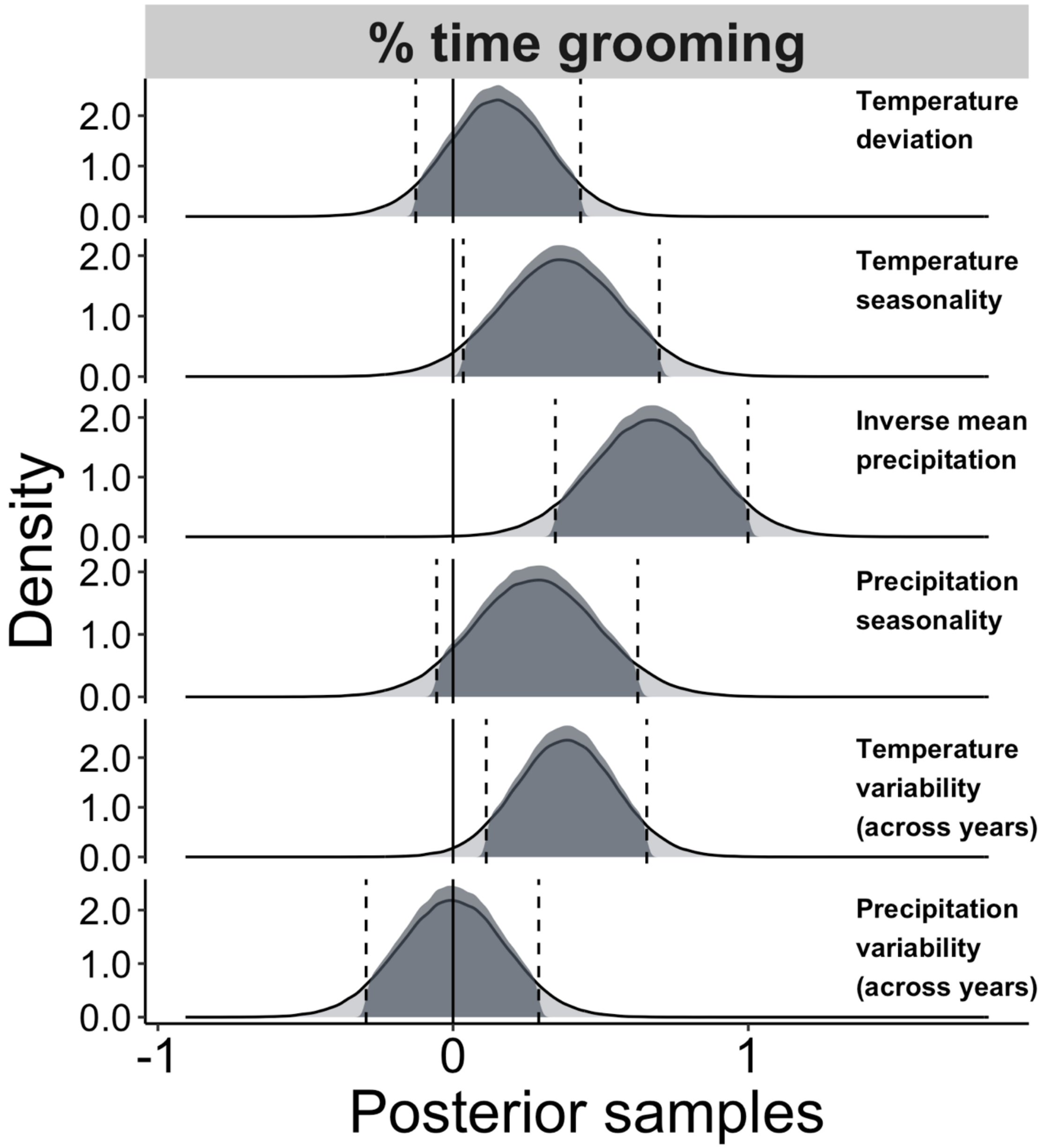


**Fig. S3:** Density of posterior samples of the four different climate variables used to calculate the ICC along with temperature and precipitation variability across years when modeled as individual predictors of percent time spent grooming. Each cell contains the density of posterior samples of a climate variable for percent time spent grooming across retained iterations from 100 MCMCglmm models run with 100 randomly sampled phylogenies from Upham et al. (2019). The dashed lines and shading represent the 89% credible intervals for these samples.

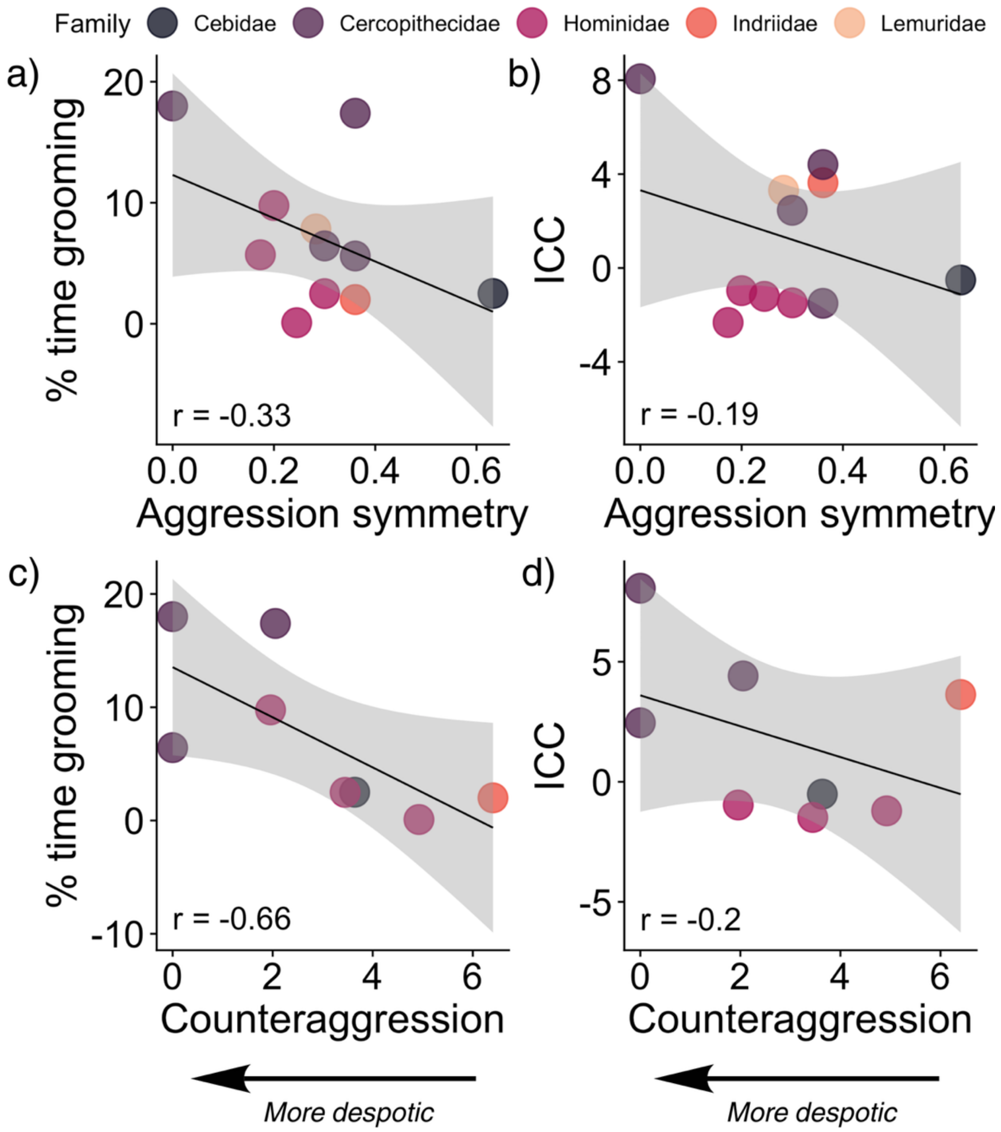


**Fig. S4:** More despotic species groom more and score higher on the ICC. Correlations between mean percent time grooming and a) aggression symmetry and c) counteraggression, and ICC and b) aggression symmetry and d) counteraggression. The square-root of aggression symmetry and counteraggression were used for the purposes of plotting, while correlation coefficients come from raw values presented in Kavanagh et al. (2021).

### SUPPLEMENTARY REFERENCES

Creighton, M. J. A., & Nunn, C. L. (2023). Explaining the primate extinction crisis: predictors of extinction risk and active threats. *Proc. R. Soc. B.*, 290(2006), 20231441.

DeCasien, A. R., Williams, S. A., & Higham, J. P. (2017). Primate brain size is predicted by diet but not sociality. *Nat. Ecol. Evol.*, 1(5), 0112.

Fritz, S. A., & Purvis, A. (2010). Selectivity in mammalian extinction risk and threat types: a new measure of phylogenetic signal strength in binary traits. *Conserv. Biol.*, 24(4), 1042-1051.

Grueter, C. C., Bissonnette, A., Isler, K., & van Schaik, C. P. (2013). Grooming and group cohesion in primates: implications for the evolution of language. *Evol. Hum. Behav.*, 34(1), 61-68.

Herrera, J. P., Moody, J., & Nunn, C. L. (2023). Predicting primate–parasite associations using exponential random graph models. *J. Anim. Ecol.*, 92(3), 710-722.

Kavanagh, E., Street, S. E., Angwela, F. O., Bergman, T. J., Blaszczyk, M. B., Bolt, L. M., *et al*. (2021). Dominance style is a key predictor of vocal use and evolution across nonhuman primates. *R. Soc. Open Sci.*, 8(7), 210873.

Pagel, M. (1999). Inferring the historical patterns of biological evolution. *Nature*, 401(6756), 877-884.

Revell, L. J. (2024). *phytools: Phylogenetic Tools for Comparative Biology (and Other Things) (Version 2.5-2)*. R package. Available at: https://cran.r-project.org/package=phytools.

Upham, N. S., Esselstyn, J. A., & Jetz, W. (2019). Inferring the mammal tree: species-level sets of phylogenies for questions in ecology, evolution, and conservation. *PLoS Biol.*, 17(12), e3000494.
